# Uncovering the drivers of host-associated microbiota with joint species distribution modeling

**DOI:** 10.1101/137943

**Authors:** Johannes R. Björk, Francis K.C. Hui, Robert B. O’Hara, Jose M. Montoya

## Abstract

In addition to the processes structuring free-living communities, host-associated microbiota are directly or indirectly shaped by the host. Therefore, microbiota data have a hierarchical structure where samples are nested under one or several variables representing host-specific factors, often spanning multiple levels of biological organization. Current statistical methods do not accommodate this hierarchical data structure, and therefore cannot explicitly account for the effect of the host in structuring the microbiota. We introduce a novel extension of joint species distribution models (JSDMs) which can straightforwardly accommodate and discern between effects such as host phylogeny and traits, recorded covariates like diet and collection sites, among other ecological processes. Our proposed methodology includes powerful yet familiar outputs seen in community ecology overall, including: (i) model-based ordination to visualize and quantify the main patterns in the data; (ii) variance partitioning to asses how influential the included host-specific factors are in structuring the microbiota; and (iii) co-occurrence networks to visualize microbe-to-microbe associations.

## Introduction

Ecological communities are the product of stochastic and deterministic processes; while environmental factors may set the upper bound on carrying capacity, competitive and facilitative interactions within and among taxa determine the identity of the species present in local communities. Ecologists are often interested in inferring ecological processes from patterns and determining their relative importance for the community under study [1]. During the last few years, there has been a growing interest in developing new statistical tools aimed toward ecologists and the analysis of multivariate community data (see e.g., [2]). Many of the distance-based approaches however, have a number of drawbacks, including uncertainty of selecting the most appropriate null models, low statistical power, and the lack of possibilities for making predictions [3]. One alternative, model-based framework which has become increasingly popular in community ecology is joint species distribution models (JSDMs, [4]). Such models are an extension of generalized linear mixed models (GLMMs, [5]), where multiple species are analyzed simultaneously often together with environmental variables, thereby revealing community level responses to environmental change. By incorporating both fixed and random effects, sometimes at multiple levels of biological organization, JSDMs have the capacity to assess the relative importance of processes such as environmental and biotic filtering versus stochastic variability. Furthermore, with the increase of trait-based and phylogenetic data in community ecology together with the growing appreciation that species interactions are constrained by the “phylogenetic baggage” they inherit from their ancestors [6], JSDMs can further accommodate information on both species traits and phylogenetic relatedness among species [7,8,9,10]. Finally, accounting for phylogenetic relatedness among species can greatly improve estimation accuracy and power when there is a phylogenetic signal in species traits and/or residual variation ([11]).

To model covariances between a large number of species using a standard multivariate random effect, as a standard JSDM [4, 12] does, is computationally challenging; the number of parameters that needs to be estimated when assuming a completely unstructured covariance matrix increases rapidly (quadratically) with the number of species. An increasingly popular tool for overcoming this problem, which is capable of modeling such high-dimensional data, is latent factor models [13]. In community ecology, latent factor models and JSDMs have been combined to allow for a more parsimonious yet flexible way of modeling species covariances in large communities [10, 14]. Such an approach offers a number of benefits. First, latent factors provide a method of explicitly accounting for residual correlation. This is important because missing covariates, ecological interactions and/or spatio-temporal correlation will induce residual correlation among species, which, if not accounted for, may lead to erroneous inference. Second, latent factors facilitate model-based ordination in order to visualize and quantify the main patterns in rows and/or columns of the data [15, 16]. While traditional distance-based ordination techniques may confound location (i.e., the mean abundance) and dispersion (i.e., the variability) effects [3], model-based ordination directly models the mean-variance relationship and can therefore accurately distinguish between the two effects [17, 18]. Finally, the estimated factor loadings can be conveniently interpreted as indicating whether two species co-occur more or less often than by chance as well as the direction and strength of their co-occurrence, thus allowing a latent factor approach to robustly estimate large species-to-species co-occurrence networks [19]. Note that an important decision when fitting latent factor models, is the choice of the number of latent factors. While less than five is usually sufficient for a good approximation to correlations, there is a trade-off between model complexity and the model’s capacity to capture the true correlation structure ([13]). An alternative approach is to use variable selection, which automatically shrinks less-informative latent factors to zero ([20]).

In parallel to community ecology, there is a growing field of microbial ecology studying both free-living and host-associated microbiota. While microbial ecologists can adopt many of the same statistical tools developed for traditional multivariate abundance data (see e.g., [21]), researchers studying host-associated microbiota need to consider an additional layer of processes structuring the focal community, namely that host-associated microbiota are addition ally shaped directly or indirectly by their hosts. For example, interactions between hosts and microbes often involve long-lasting and sometimes extremely intimate relationships, where the host may have evolved the capacity to directly control the identity and/or abundance of its microbial symbionts [22, 23]. Similar to an environmental niche, the host must be viewed as a multidimensional composite of all host-specific factors driving the occurrence and/or abundance of microbes within a host–everything from broad evolutionary relationships between host species [24] to the direct production of specific biomolecules within a single host individual [25]. As a result, host-associated microbiota have a hierarchical data structure where samples are nested under one or several variables representing recorded and/or measured host-specific factors sometimes spanning multiple levels of biological organization.

In this article, we propose a novel extension of JSDMs to analyze host-associated microbiota, based around explicitly modeling its characteristic hierarchical data structure. In doing so, our proposed model can straightforwardly accommodate and discriminate among any measured host-specific factors. Over the past few years, there has been an increase of model-based approaches aimed specifically toward the analysis of host-associated microbiota (see e.g., [12, 26, 27, 28]). To our knowledge however, our proposed model is the first to explicitly and transparently account for the aforementioned hierarchical structure that is inherent in data on host-associated microbiota (Fig 1). Other key features of the proposed model, which are inherited from JSDMs and latent factor models, include: (1) parsimonious modeling of the high-dimensional correlation structures typical of host-associated microbiota; (2) model-based ordination to visualize and quantify the main patterns in the data; (3) variance partitioning to assess the explanatory power of the modeled host-specific factors and their influence in shaping the microbiota; and finally (4) co-occurrence networks to visualize OTU-to-OTU associations. Furthermore, by building our model in a probabilistic, i.e., Bayesian framework, we can straightforwardly sample from the posterior probability distribution of the correlation matrix computed by the factor loadings; this means that we can choose to look at, or further analyze the correlations that have at least e.g., 95% (or even 97% or 99%) probability.

**Figure 1:**
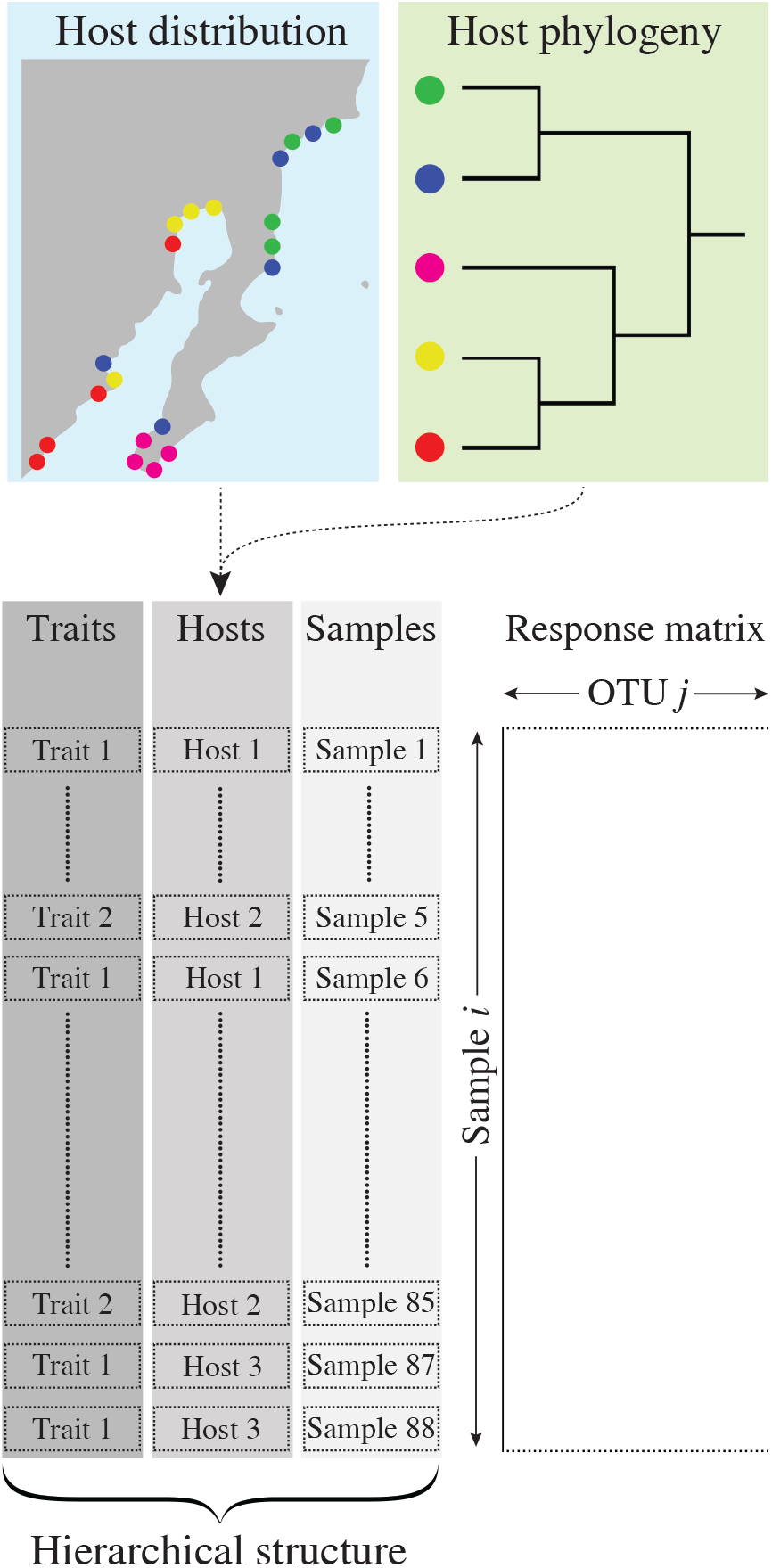
Host-associated microbiota data have a hierarchical data structure. In this example, samples are nested within host species which in turn are nested under species traits. As there are also data on the host’s geographical distribution, host species can be further nested within observation/collection sites. Additional data that are often available is the host species phylogeny. The proposed model extension can straightforwardly accommodate for this hierarchical data structure and discriminate their importance in structuring the microbiota.

We apply our proposed model to two published data sets. While we include the effect of host phylogenetic relatedness in both case studies, we illustrate the flexibility of our approach by adapting the proposed model to overdispersed counts and presence-absence responses, and study-specific meta data relevant to each case study. By utilizing recent progress in latent factor modeling, our proposed model can also assist in cases where meta data are scarce by finding latent “hidden” variables driving the microbiota.

## Methods

We applied the proposed methodology to two published data sets on host-associated microbiota. Both datasets possess two main features which characterize many host-associated microbiota data, namely high dimensionality i.e., the number of OTUs is a non-negligable proportion of the number of samples, and sparsity i.e., most OTUs are rarely observed. The first data set comprise 90 samples from 20 sponge species collected in four closely located sites in the Bocas del Toro archipelago (Fig S1, for original study see [29]). The meta data contain apart from collection site, a classification of hosts into either High Microbial Abundance (HMA) or Low Microbial Abundance (LMA) sponges (hereafter termed *ecotype*). This classification is based on the abundance of microbes harbored by the host and determined by transmission electron microscopy [30]. The authors constructed a host phylogeny from 18S rRNA gene sequences (downloaded from GenBank) by implementing a relaxed-clock model in MrBayes. The data have a hierarchical structure with *n* = 90 samples nested within *S* = 20 host species and *L* = 4 collection sites. Host species are then further nested under one of *R* = 2 ecotypes. The response matrix had already been filtered to only include OTUs (defined at 97% similarity) with at least 500 reads, but we further removed OTUs with less than 20 presences across samples, resulting in *m* = 187 modeled OTUs.

The second data set consists of 59 neotropical bird species with a total of 116 samples from the large intestine. Host species were collected from 12 lowland forests sites across Costa Rica and Peru (Fig S2, for original study see [31]). The meta data include bird taxonomy and several covariates–including dietary specialization, stomach contents and host habitat. The authors sequenced and used the mitochondrial locus ND2 to reconstruct the host phylogeny by implementing a partitioned GTR + Γ model in BEAST. Similarly to the sponge data set, this data set has a hierarchical structure with *n* = 116 samples nested within *S* =59 host species and *L* = 12 collection sites. We filtered the response matrix to include OTUs (defined at 97% similarity) with at least 50 reads and 40 presences across samples, resulting in *m* = 151 modeled OTUs. Of the full list of covariates available, we included diet, stomach content, sex, elevation and collection site as explanatory predictor variables in our model. While diet and geography have been shown to influence the human gut microbiota (see e.g., [32, 33]), the effect of sex and elevation is less known.

## Joint species distribution models

We considered two response types commonly encountered in host-associated microbiota data: counts and presence-absence. Formally, let the response matrix being modeled consist of either counts or presence-absence records of *m* OTUs from *n* samples, and let *y_ij_* denote the response of the *j*-th OTU in the *i*-th sample. Also, let 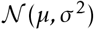 denote a univariate normal distribution with mean *μ* and variance *σ*^2^, and analogously, let 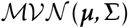 denote a multivariate normal distribution with mean vector and covariance matrix Σ. We now split our model formulation up into the two case studies/response types.

### Case Study 1 (Counts)

Due to the presence of overdispersion that was quadratic in nature, as confirmed by a mean-variance plot of the OTU counts (not shown), we assumed a negative binomial distribution for the responses. Specifically, we considered a negative binomial distribution with a quadratic mean-variance relationship for the element *y_ij_*, such that 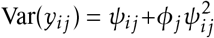 where *φ_j_* is the OTU-specific overdispersion parameter. The mean abundance was related to the covariates using a log-link function. Denoting the mean abundance of OTU *j* in sample *i* by *ψ_ij_*, then we have

**Model 1**

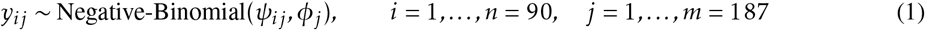

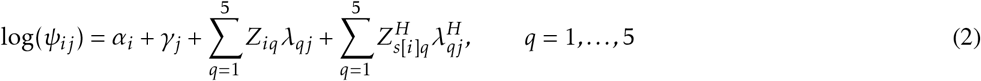

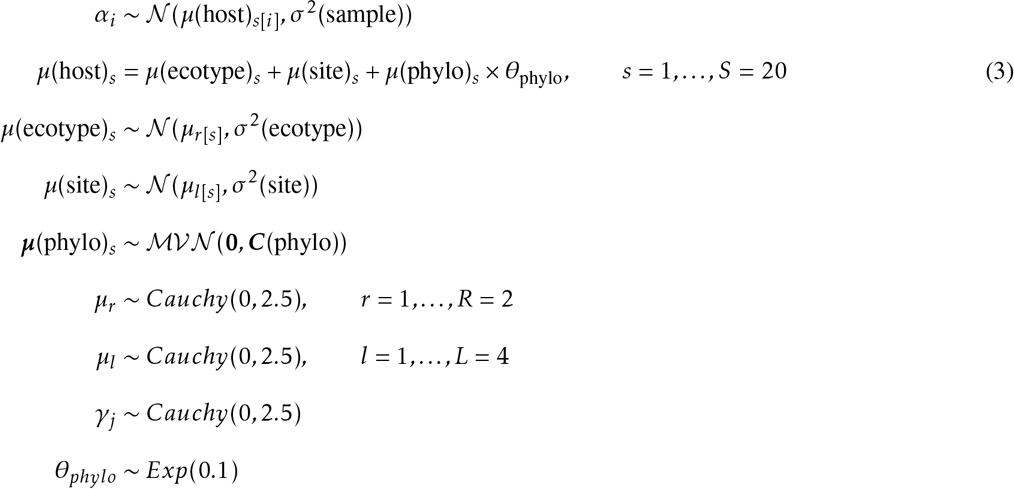

To clarify the above formulation, *s, r* and *l* index effects that are attributed to the *S* = 20 host species, *R* = 2 ecotypes and *L* = 4 sites respectively. For instance, “*s*[*i*]” and “*r*[*s*]” denote “sample *i* nested within host species *s*” and “host species *s* nested within ecotype *r*”, respectively (Fig 1). In equation (2), the quantities *α_i_* and *γ_j_* represent sample and OTU-specific effects, respectively. The former adjusts for differences in sequencing depth among samples, while the latter controls for differences in OTU total abundance. The inclusion of *α_i_* serves two main purposes. First and foremost, including *α_i_* allows us to account for the hierarchical data structure and its effect on sample total abundance specifically. In particular, to account for sample *i* being nested within host species *s* (which are further nested within ecotype *r*) and site *l*, the sample effects *α_i_* are drawn from a normal distribution with a mean that is a linear function of three host-specific effects: host ecotype *μ*(ecotype); host collection site *μ*(site); and host phylogeny *μ*(phylo). Furthermore, the host ecotype *μ*(ecotype) and host collection site *μ*(site) effects are themselves drawn from a normal distribution with an ecotype and site-specific mean, respectively. Second, the inclusion of *α_i_* means that the resulting ordinations constructed by the latent factors on the sample *Z_iq_* and host species 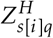 level are in terms of species composition only, as opposed to a composite of abundance and composition if the site effects were not included in the formulation. We included five latent factors at both the sample and host species level, and both *Z_iq_* and 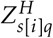 were assigned standard normal priors 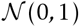 with the assumption of zero mean and unit variance to fix the location and scale (see Chapter 5, [34]). Furthermore, to address rotational variance, the upper triangular component of both loading matrices (i.e., sample ***λ*** and host species ***λ***^*H*^ level) are fixed to zero with the diagonals constrained to be positive [35]. As recommended by Polson and Scott [36], and analogous to the prior distributions we use for the mean *μ_r_* and *μ_l_*, we used a weakly informative prior in the form of a half-Cauchy distribution with a center and scale equal to 0 and 2.5 for the overdispersion parameter *ψ*. Moreover, following Gelman et al. [37], we used the same distribution with location and scale equal to 0 and 1 as prior information on the variance parameters: *σ*^2^(sample); *σ*^2^(ecotype); and *σ*^2^(site). Based on our empirical investigation, we found that the use of such priors stabilized the MCMC sampling substantially without introducing too much prior information, compared to using more uninformative prior distributions. Lastly, the quantity ***C***(phylo) corresponds to a phylogenetic correlation matrix constructed from the host phylogeny by assuming Brownian motion evolution such that the covariances between host species are proportional to their shared branch length from the most recent common ancestor [38]. The phylogenetic parameter *θ*_phylo_ quantifies variance that can be attributed to the phylogenetic effect, and is drawn from an exponential distribution with a rate parameter of 0.1. Similar to the half-Cauchy priors, this prior distribution provides a weak level of regularization–a rate parameter of 0.1 gives a prior mean of 10, thus preventing the estimated variance of getting implausibly large.

### Case Study 2 (Presence-absence)

We modelled the presence (*y_ij_* = 1) or absence (*y_ij_* = 0) of OTU *j* in sample *i* using probit regression, implemented via the indicator function 1*_z_ij__*>0 where the latent score is normally distributed with the mean equal to a linear function of the covariates and latent factors, and variance set equal to one. The hierarchical model was set up as follows:

**Model 2**

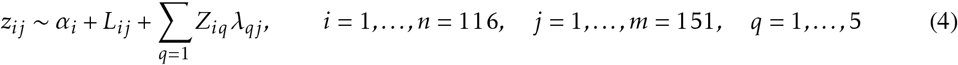

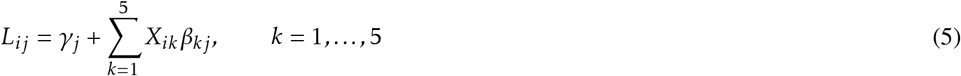

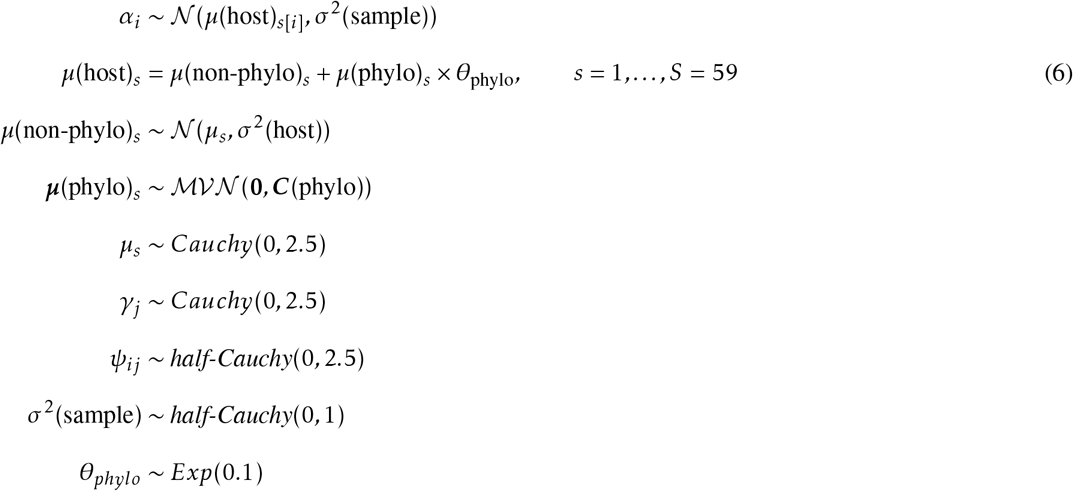

While the above description is largely the same as that of *Model 1*, we also included here a linear predictor *L_ij_* to model the effects of five available covariates (represented by the model matrix *X_ik_; k* = 1,…, 5) on species composition (equation (5)). The linear predictor *L_ij_* thus acts to explain covariation between OTUs due to the measured explanatory predictor variables, while the latent factors account for the remaining, residual covariation. Similarly to *Model 1*, including *α_i_* means that the covariation between OTUs is in terms of species composition only. By drawing the sample effects *α_i_* from a normal distribution with a mean that is a linear function of both non-phylogentic *μ*(*non – phylo*) and phylogenetic *μ*(phylo) host effects (equation (6)), we account for the hierarchical structure present in the data. Furthermore, from the loading matrix λ, we computed a covariance matrix as Ω = ***λλ***^⊤^, which we subsequently convert to a correlation matrix for studying the OTU-to-OTU co-occurrence network.

For both case studies, we used Markov Chain Monte Carlo (MCMC) to estimate the models `via JAGS` [39] and the `run jags` package [40] in `R` [41]. For each model, we ran one chain with dispersed initial values for 300,000 iterations saving every 10^*th*^ sample and discarding the first 25% of samples as burn-in. We evaluated convergence of model parameters by visually inspecting trace and density plots using the R packages *coda* [42] and *mcmcplots* [43], as well as using the Geweke diagnostic [44].

### Variance partitioning

To discriminate among the relative contributions of the various factors driving covariation in the JSDMs, we partition the explained variance by the row effects (*α_i_*), the linear predictor (*L_ij_*), and the loadings (*λ_qj_* and 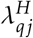) into components reflecting sample and host level effects. Such a variance decomposition is analogous to the sum-of-squares and variance decompositions seen in Analysis of Variance (ANOVA) and linear mixed models ([45]). Depending on the response type, the row effects capture variance in relative abundance (*Model 1*) or species richness (*Model 2*), while the linear predictor and the loadings capture variance in species composition. As mention above, when the linear predictor is included in (*Model 2*), the loadings capture residual variation not accounted for by the modeled covariates. Variance partitioning therefore allows us to asses the explanatory power of the hierarchical data structure, and measured covariates including “hidden” factors, and how influential each of them are in structuring the host-associated microbiota ([10]).

We now discuss in more detail how we partition the explained variance into components attributed to the row effects (*α_i_*) for *Model 1*, and the loadings (*λ_qj_*) together with the linear predictor (*L_ij_*) for *Model 2*. Let *V*_total_ denote the total variance of the *α_i_*, while *V*_sample_, *V*_ecotype_, *V*_site_ and *V*_phylo_ denote the variances for the sample, host ecotype, host collection site and host phylogeny, respectively. Then for Case Study 1 we have,

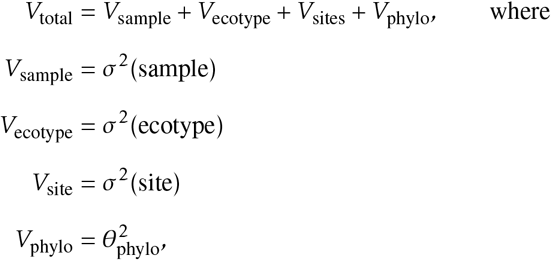

and for Case Study 2 we have,

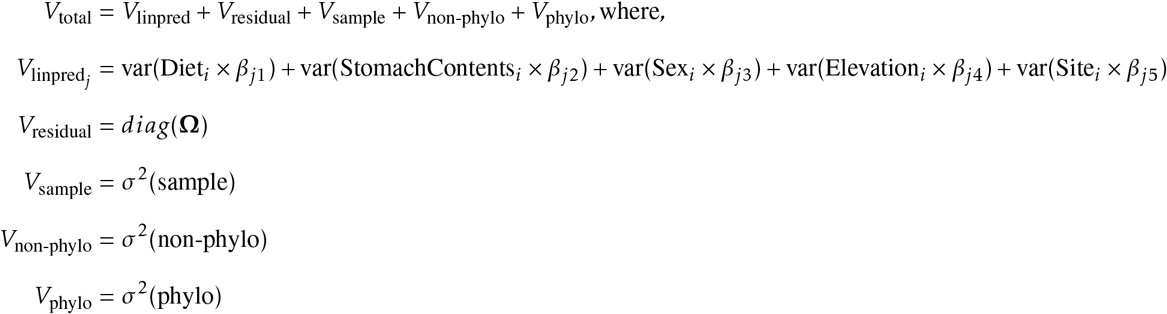

In the second partitioning, the quantity *V*_linpred_ represents the variance explained by the linear predictor *L_ij_*, the *V*_residual_ represents the residual variance not accounted for by the modeled predictor variables i.e., as explained by the diagonal elements of the residual covariance matrix Ω, and finally the *V*_sample_, *V*_non-phylo_ and *V*_phylo_ to variance attributed to the hierarchy present on the row effects *α_ij_*.

## Results

Below we present the main results for each case study. We used the 95% highest density interval (HDI) as a measure of statistical significance. That is, if a parameter or a pairwise parameter comparison excludes zero, then we conclude that the posterior probability of the difference being significantly different from zero exceeds 95%.

### Case study 1

We applied *Model 1* to data on sponge host-associated microbiota [29]. The fitted model revealed that more than 86% of the variation in relative abundance among samples could be attributed to processes operating on the host-species level (Table 1; Fig 2). More specifically, 57% of this variation was explained by host phylogenetic relatedness, even though the 95% HDI for the phylogenetic effects did not exclude zero for any of the host species. While this suggests the presence of a phylogenetic signal in one or more host traits affecting microbial abundance and/or occurrence, it also indicates that no particular host species or host species clade have a stronger signal than the rest. Easson and Thacker [29] used the Bloomberg’s K statistic and found a significant signal of the host phylogeny on the inverse Simpson’s index. This index measures the diversity of a community, but is strongly influenced by the relative abundance of its most common species ([46]). The authors specifically noted that host species *Aiolochroia crassa, Aplysina cauliformis* and *Aplysina fulva* from the order Verongida, along with host *Erylus formosus* from the order Astrophorida had higher values of this index compared to the rest of the host species. Similarly, we found that the same four hosts harbored more abundant (Fig 2) and distinctively different microbiotas than the other host species (Fig 3). Pairwise comparisons of these four hosts showed that *A. crassa* harbored markedly different microbial composition compared to its two closest relatives *A. cauliformis* and *A.fulva* (Table S1; Table S2). These three hosts were nonetheless collected at the same site. The two species from the genus *Aplysina* on the other hand, harbored very similar microbiota composition to that of host *E. formosus* even if they were collected some 17,000 km apart.

**Figure 2:**
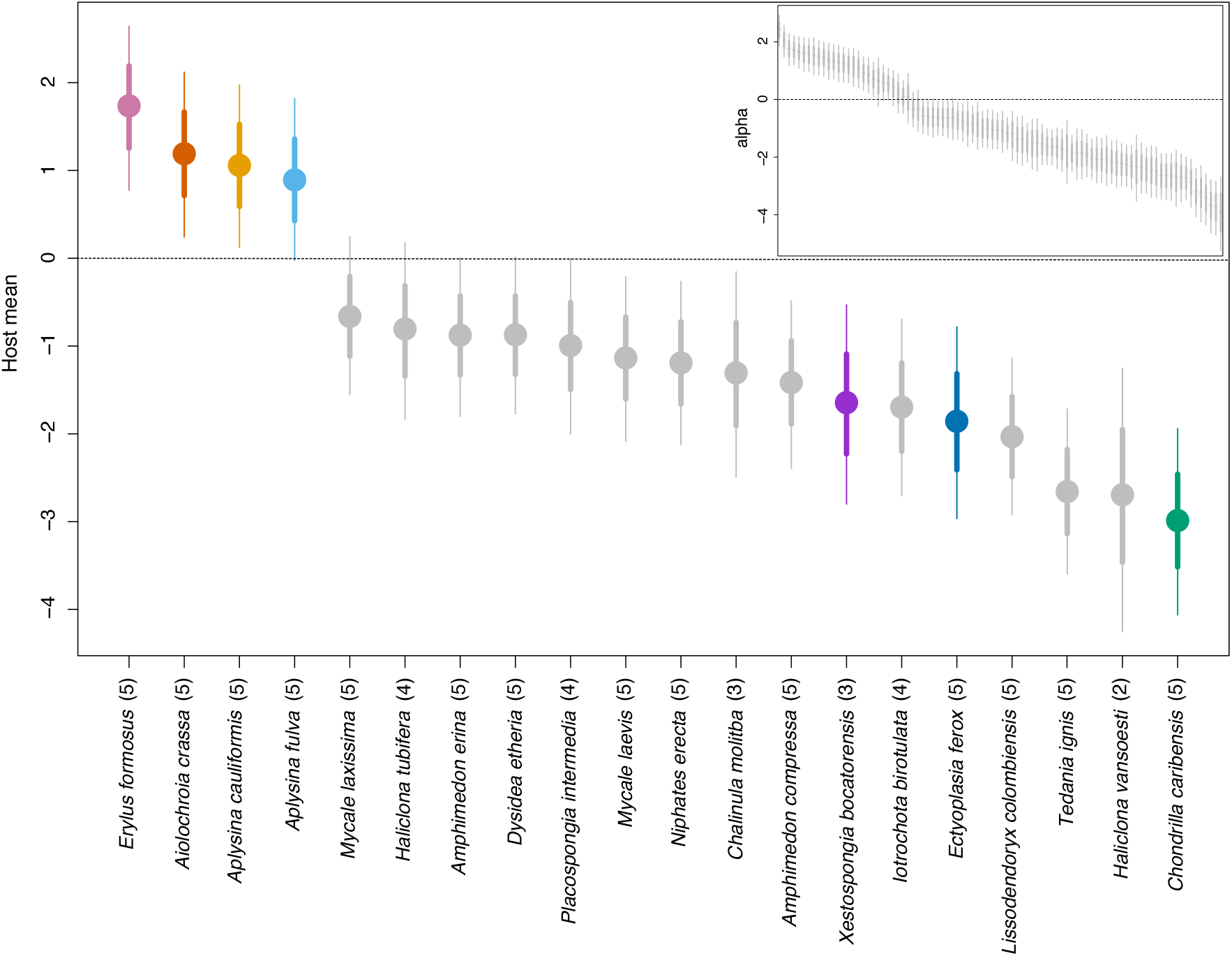
The main plot shows a caterpillar for the host means *μ*(host)_*s*_, with the colors representing the 7 HMA hosts. The subplot shows a caterpillar plot for the row effects *alpha_i_*. The quantiles corresponds to the 95% (thin lines) and 68% (thick lines) credible intervals. The number within the parentheses indicates how many individuals per host species were used to draw inference on.

**Figure 3:**
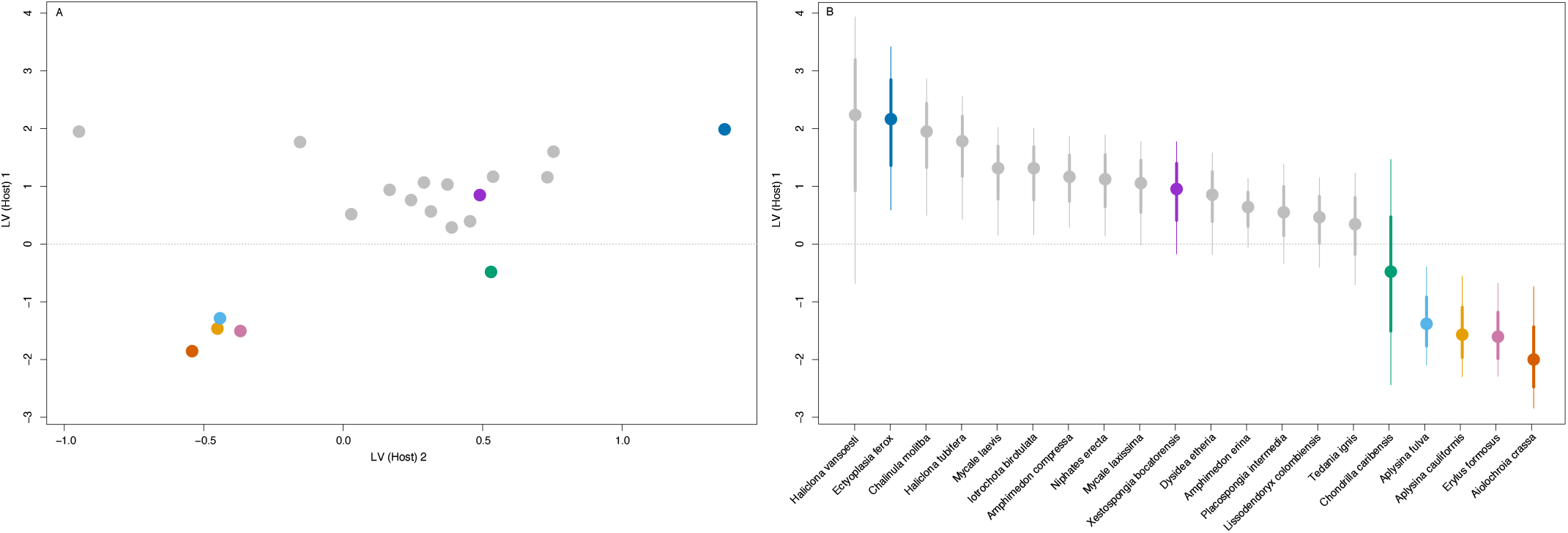
The left plot (A) shows the ordination constructed by the latent factors on the host species level ***Z^H^***, and the right plot (B) shows the corresponding caterpillar for first latent factor 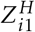. The quantiles corresponds to the 95% (thin lines) and 68% (thick lines) credible intervals.

**Figure 4:**
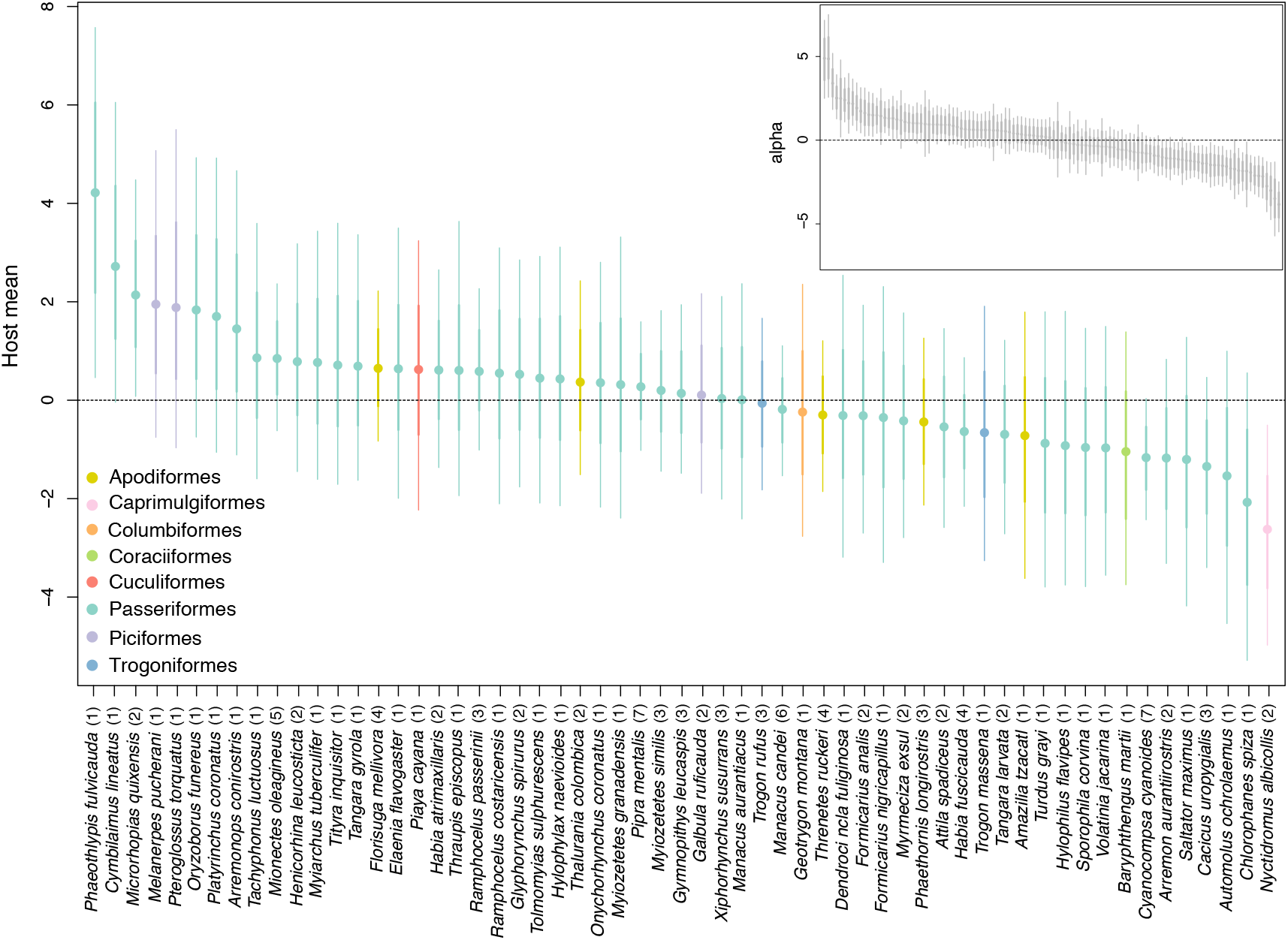
The main plot shows a caterpillar for the host means *μ*(host)_*s*_ colored by host taxonomy at the order level, while the subplot shows a caterpillar plot for the row effects *alpha_i_*. The quantiles corresponds to the 95% (thin lines) and 68% (thick lines) credible intervals. The number within the parentheses indicates how many individuals per host species were used to draw inference on.

**Table 1:** Variation explained by the hierarchy present on *alpha_i_*, i.e., the host effects *μ*(host)_*s*_.

|  |  |
| --- | --- |
| Phylogeny | 57.09% |
| Ecotype | 14.58% |
| Site | 14.51% |
| Sample | 13.82% |

Host ecotype and collection site roughly explained two thirds of the remaining variation in relative abundance (Table 1). Furthermore, the host species level explained 39% of the variation beyond differences in relative abundance, with the remaining variation explained by the latent factors on the sample level. While samples did not cluster based on ecotype or sites, samples belonging to HMA hosts generally formed tighter clusters compared to samples from LMA hosts (Fig S3). Note however that because the sampling scheme in the original study confounded host ecotype and collection site, it is impossible to fully disentangle the two.

### Case study 2

Fitting *Model 2* to the data on neotropical bird gut-associated microbiota [31] revealed that only 9% of the variation in species richness among samples could be explained by processes acting on the host species level, including processes related to the host phylogeny. The remaining 91% of this variation was captured by processes operating on the sample level (Table 2). Of the total variance in species occurrence, variation in species richness only accounted for, on average, about 17%. The modeled predictor variables explained 69% of the total variance, and varied from a minimum of less than 0.01% to a maximum of 99.7% across all OTUs (Fig 5). The predictor variable that had the largest average effect on microbiota composition was collection site (21.33%, Table 2). None of the estimated regression coefficients for the predictor variables excluded zero (Fig S4). Furthermore, the ordination plots constructed from the the first two latent factors did not reveal any obvious clustering by e.g., host taxonomy (at the order level), collection site, or diet (broad dietary specialization) (Fig 6; Fig S5; Fig S6).

**Figure 5:**
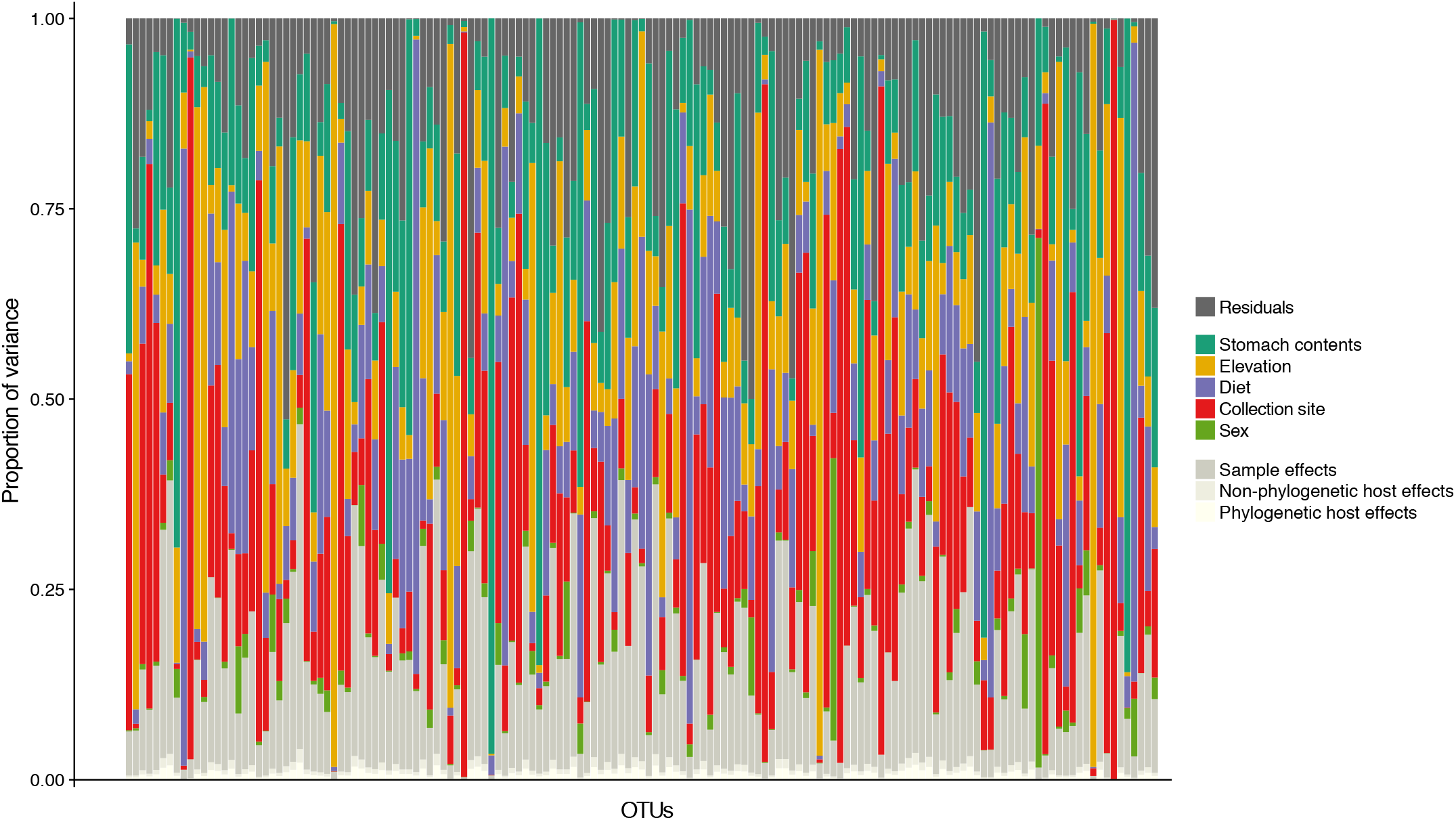
The y-axis shows the relative proportion of variance in species occurrences explained by the hierarchy present on *alpha_i_*, the covariates included on the linear predictor *L_ij_*, and the residual variance not accounted for by the modeled effects i.e., the diagonal elements of the residual covariance matrix **Ω**, for each OTU (x-axis).

**Figure 6:**
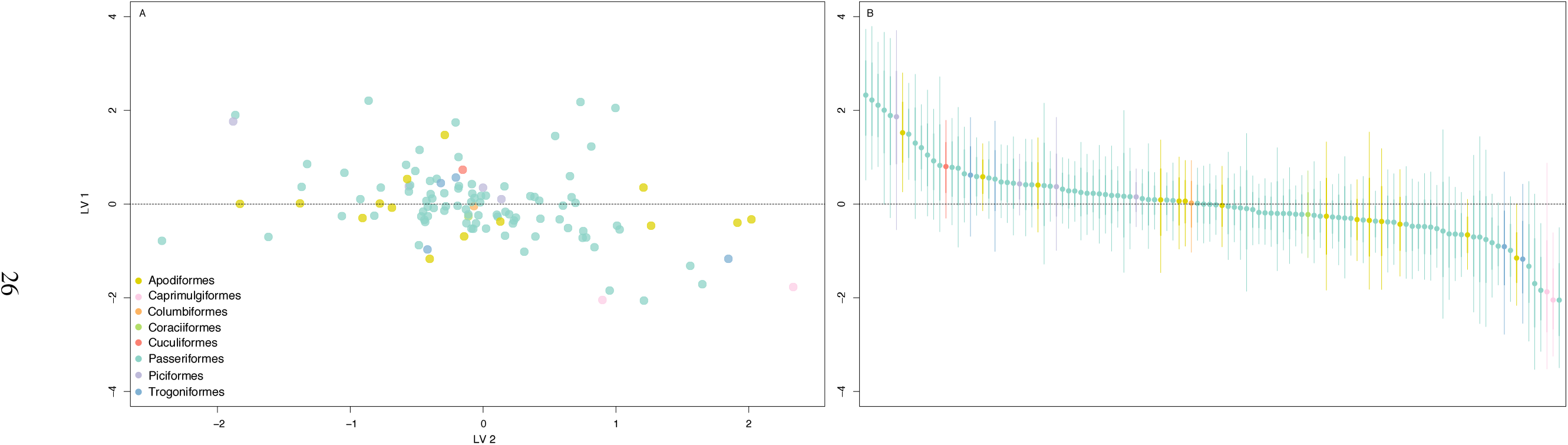
The left plot (A) shows the ordination constructed by the latent factors **Z** colored by host taxonomy (at the order level), and the right plot (B) shows the corresponding caterpillar for first latent factor *Z*_*i*1_. The quantiles corresponds to the 95% (thin lines) and 68% (thick lines) credible intervals.

**Table 2:** Variation attributed to the linear predictor *L_ij_*, the residual variation captured the diagonal elements of the residual covariance matrix **Ω**, and by the hierarchy present on the row effects *α_ij_*, i.e., the host effects *μ*(host)_*s*_.

|  |  |
| --- | --- |
| Collection site | 21.33% |
| Stomach contents | 16.13% |
| Elevation | 15.97% |
| Diet | 13.59% |
| Sex | 2.12% |
| Residuals | 13.89% |
| Sample | 15.5% |
| Non-Phylogeny | 0.65% |
| Phylogeny | 0.82% |

We ran an edge betweeness community detection algorithm [47] on the correlation matrix computed from the loading matrix ***λ*** where links represent positive and negative co-occurrences with at least 95% posterior probability. We colored nodes by their bacterial taxonomic affiliation at the phylum level. This revealed a large tightly knit cluster with well connected nodes in the centre and less connected nodes in the periphery of the cluster. The network displayed equal proportion of positive and negative co-occurrences, and with no apparent clustering of OTUs belonging to certain phyla (Fig S7). Caution should, however, be taken when interpreting statistical interactions: these are residual species-to-species co-occurrences that can only be considered as hypotheses for ecological interactions, and without additional biological information it is impossible to definitively confirm or assess their nature ([19, 48, 49]).

## Discussion

In this paper, we have developed a joint species distribution model (JSDM) aimed towards analyzing host-associated microbiota data. The present work builds upon and extends existing JSDMs by specifically targeting the hierarchical structure implicit in host-associated microbiota studies, while also including several other features that are attractive for analyzing such data. First, we have shown how overdispersed counts and presence-absence data, two common features of host-microbiota data can be modeled under a single framework by implementing a negative binomial and a probit distribution with the appropriate link function. Furthermore, we have utilized recent progress in latent factor modeling in order to represent the high-dimensional nature of host-microbiota data as a rank-reduced covariance matrix, thus making the estimation of large OTU-to-OTU covariance matrices computationally tractable. By doing so, we have also demonstrated how latent factors, both alone or together with measured covariates, can be used for variance partitioning and further visualized as ordinations and co-occurrence networks. Lastly, depending on the modelled response function, we have illustrated that the variance partitioning of the hierarchy present on the rows can be represented in terms of either relative abundance or species richness.

We adapted our proposed model to make use of two published data sets on host-associated microbiota. Although our goal was not to compare the results from these two case studies, such a systematic comparison can be done using a model-based approach like ours. Broadly, the data analyzed here suggest that markedly different processes are shaping the microbiota harbored by these different host organisms. Individually, the main results from each of our two models were generally in agreement with the results reported in their respective original study; for example, *Model 1* identified the same four host species reported by Easson and Thacker [29] to have more abundant and distinctively different microbiotas compared to the other analyzed hosts. Similarly to Hird et al. [31], the ordinations produced by *Model 2* did not cluster by host diet, host taxonomy nor collection site. By partitioning variance among fixed and random effects, *Model 2* further showed that there was substantial variation across OTUs in terms of which predictor variables explained the most variance.

While distance-based methods such as PERMANOVA still remains one of the most widely used non-parametric methods to analyze host-associated microbiota data, model-based approaches are increasingly recognized to outper form such analyses (see e.g., [3, 17, 27]), and we see our proposed model as making a strong case for further empirical comparisons between distanced-based and model-based approaches to analyzing microbiota data.

There are a number of extensions one could make to the proposed model. Perhaps the most important of these stems from the growing recognition that high-throughput DNA sequencing produces compositional data, i.e., non-negative counts with an arbitrary sum imposed by the sequencing platform, which can produce spurious correlations if not properly accounted for (see e.g., [50, 51, 52]). Because of the log-link function used in *Model 1*, it is possible to parameterize this model and regard it in terms of compositional effects (see [53] and also noting the fact that the negative binomial distribution can itself be parameterized as a hierarchical Poisson model with Gamma distributed random effects), although for ease of estimation and interpretation we chose to adopt the standard negative binomial parameterization. This topic remains an area of active research, and there are currently several model-based methods (see e.g., [54, 55, 56, 57]) to infer co-occurrence networks, each with its own set of assumptions–it is not yet conclusive that any one of these methods outperforms the rest. Other model extensions and modifications can also be made in order to answer specific ecological questions of interest. For example, whether closely related host species harbor closely related microbes (i.e., host-microbiota phylogenetic congruence), or whether similarity among host-associated microbiota decreases as a function of increasing geographical distance or social connectance between hosts. Such questions may be answered for instance, by incorporating a phylogenetic effect acting on the columns of the response matrix, and by implementing a Gaussian process model that quantifies the degree of spatial and/or social autocorrelation between hosts, respectively. These two “flavors” of JSDMs and mixed models more generally have previously been considered in community ecology, both separately [58, 59, 60] and combined [8], although both computation and successful estimation and inference of all the model parameters remain a major issue especially with the high-dimensional nature of host-associated microbiota data. In summary, while substantial methodological advances have been made over the past few years in developing an extensive model framework for community ecological data, to date there exists no similar unifying framework for modeling host-associated microbiota which is directly tailored to the hierarchical and correlation structures present as well as questions of interest specific to such data. Our proposed model, which explicitly accounts for the host’s effect in structuring its microbiota, takes us closer to that goal.

## Acknowledgements

We thank Dr. Robert W. Thacker and Dr. Sarah Hird for sharing their data sets on the microbiota associated to marine sponges and neotropical bird species, respectively. We further thank three anonymous reviewers for providing constructive feedback and comments. J.R.B was supported by an FPI Fellowship from the Spanish Government (BES-2011-049043). J.M.M. was supported by the French LabEx TULIP (ANR-10-LABX-41; ANR-11-IDEX-002-02), by the Region Midi-Pyrenees project (CNRS 121090) and by the FRAGCLIM Consolidator Grant, funded by the European Research Council under the European Union’s Horizon 2020 research and innovation programme (grant agreement number 726176)

## Code and Data Availability

All code and data are available on Open Science Framework (`DOI:10.17605/OSF.IO/T9NXH`) with a tutorial on how to fit the model and analyze the output. While we used `JAGS` to fit the model in this study, we have translated the model into `Greta` ([61]) which is an `R` style probabilistic language that scales better to large data sets.

